# Phenotypical microRNA screen reveals a noncanonical role of CDK2 in regulating neutrophil migration

**DOI:** 10.1101/574335

**Authors:** A.Y. Hsu, D. Wang, S. Liu, J. Lu, R. Syahirah, D.A. Bennin, A. Huttenlocher, D.M. Umulis, J. Wan, Q. Deng

## Abstract

Neutrophil migration is essential for inflammatory responses to kill pathogens, however it also causes tissue injury. To discover novel therapeutic targets that modulate neutrophil migration, we performed a neutrophil-specific microRNA overexpression screen in zebrafish, and identified eight microRNAs as potent suppressors of neutrophil migration. Among those, *miR-199* decreases neutrophil chemotaxis in zebrafish and human neutrophil-like cells. Intriguingly, in terminally differentiated neutrophils, *miR-199* alters the cell cycle-related pathways and directly suppresses *cyclin-dependent kinase 2* (*cdk2)*, whose known activity is restricted to cell cycle progression and cell differentiation. Inhibiting CDK2, but not DNA replication, disrupts cell polarity and chemotaxis of zebrafish neutrophils. Chemotaxis of primary human neutrophils are also reduced by CDK2 inhibition. Furthermore, *miR-199* overexpression or CDK2 inhibition significantly improves the outcome of lethal systemic inflammation challenges in zebrafish. Together, our results reveal previously unknown functions of *miR-199* and CDK2 in regulating neutrophil migration and provide new directions in alleviating systemic inflammation.

**One Sentence Summary:** *miR-199* directly suppresses *cdk2* expression, neutrophil chemotaxis and systemic inflammation.

## Introduction

Neutrophils are the first cells recruited to an immune stimulus stemming from infection or sterile injuries via a mixture of chemoattractant cues (*1*). In addition to eliminating pathogens, neutrophils coordinate the overall inflammation by activating and producing inflammatory signals in the tissue while modulating the activation of other immune cells which in some cases leads to adverse tissue damage. Over amplified or chronic neutrophil recruitment directly leads to autoimmune diseases including rheumatic arthritis, diabetes, neurodegenerative diseases, and cancer (*2*). Dampening neutrophil recruitment is a strategy to intervene in neutrophil-orchestrated chronic inflammation (*3*). Despite intensive research over the past several decades, clinical studies targeting neutrophil migration have been largely unsuccessful, possibly due to the prominent redundancy of adhesion receptors and chemokines (*4, 5*). Additional challenges lie in the balance of dampening detrimental inflammation while preserving immunity (*6*). Neutrophil migration is governed by spatially and temporally complex dynamic signaling networks (*7*). Immune cells use both mesenchymal migration (adhesive migration on substrate) and amoeboid migration (actin protrusion driven by weak substrate interaction) to infiltrate tissue using distinct signaling molecules (*8, 9*). Extensive research is required to understand the machinery regulating neutrophil recruitment, which will lead to the development of exogenous inhibitors for clinical use.

MicroRNAs (miRNAs) are short (20-22 nucleotides) conserved non-coding RNAs that are epigenetic regulators of the transcriptome. By binding primarily to the 3’UTRs of their target transcripts and recruiting the RNA-induced silencing complex, miRNAs down-regulate gene expression (*10*). miRNAs are generally fine-tuners that suppress gene expression at modest levels, and also master regulators that target the expression of a network of genes (*11*). Recently, miRNAs and anti-miRNAs have been used in clinical trials to treat cancer and infection (*12*). In addition, they have been used as tools to screen and identify the underlying mechanisms of diseases and cell behavior (*13, 14*). In immune cells, an overexpression screen of microRNA was performed to identify microRNAs that regulate B cell tolerance (*15*). In neutrophils, the functions of the highly expressed microRNAs *miR-223* (*16*) and *miR-142* (*17*) have been well characterized. However, other miRNAs and their targets as regulators of neutrophil recruitment remain largely unknown. The absence of this knowledge might lead to missed opportunities in harnessing microRNAs and their targets in restraining neutrophilic inflammation.

The zebrafish is a suitable model organism to study neutrophil biology. They have an evolutionarily conserved innate immune system, including the phagocytes and their signaling molecules including miRNAs (*17, 18*). The transparent property of zebrafish embryos enables noninvasive imaging of neutrophil behavior under physiological conditions (*19, 20*). Additionally, high fecundity and established genetic tools make zebrafish an ideal platform for genetic screens and drug discovery (*21*).

We have previously established a system to express individual miRNAs in zebrafish neutrophils to assess their function *in vivo* (*22*). Here we performed a miRNA overexpression screen and identified *miR-199* as a novel regulator of neutrophil chemotaxis. Through *miR-199* target analysis, we identified the canonical cell cycle related gene *cdk2* as a previously unrecognized regulator of neutrophil migration. Overexpression of *miR-199* or inhibition of *cdk2* in neutrophils improved survival upon infection and sterile inflammatory challenges. Our results were further validated in human neutrophil-like cells and primary human neutrophils, supporting the evolutionary conservation of *miR-199* and Cdk2 in vertebrae neutrophil biology. These discoveries expand our current understanding of neutrophil migration and suggests a novel strategy to manage neutrophilic inflammation.

## Results

### Identification and characterization of miRNAs that regulate neutrophil recruitment

As a first step to understand microRNA mediated gene regulation in zebrafish neutrophils, we sequenced microRNAs in zebrafish neutrophils sorted from 3 day old embryos. Consistent with previous reports in human and mice, *miR-223* and *miR-142* were among the microRNAs highly expressed in zebrafish neutrophils (fig. 1A, B and Data file S1). As a control, we sequenced microRNAs in the immotile epical keratinocytes and indeed detected a distinct profile (fig.1C, D). We next performed a functional genetic screen to identify a set of miRNAs, overexpression of which, moderates neutrophil migration (fig. S1). Forty-one microRNAs with lower expression levels compared with the whole embryo or keratinocytes were selected. Individual miRNA candidates were cloned and transiently expressed in zebrafish neutrophils, together with a fluorescent reporter gene. The miRNAs whose overexpression resulted in a > 50% reduction in neutrophil recruitment to infection sites were selected (Data file S2). We generated transgenic zebrafish lines stably expressing the eight miRNAs that passed the initial screen. To rule out potential artifacts associated with the transgene integration sites, more than two independent founders were obtained for each line. In seven out of the eight lines, we found a significant decrease in neutrophil recruitment to ear infection sites, regardless of the choice of the founder (fig. 1E). Similarly, we found that neutrophil recruitment was also significantly reduced in all eight lines in a tail fin amputation inflammation model (fig. 1F and Data file S3), demonstrating that the microRNA phenotypic screen faithfully represents microRNAs that could modulate neutrophil migration. A microRNA characterized in our previous work, *miR-722*, served as a positive control (*22*). Among the eight newly identified miRNAs, *miR-146* is a known suppressor of immune cell activation (*23*), whereas the other 7 miRNAs have not been characterized for their roles in neutrophilic inflammation or migration. The overexpression of *miR-199-3* resulted in the most robust inhibition in both models (fig. 1E-J) and therefore was chosen for further characterization.

**Fig. 1.**
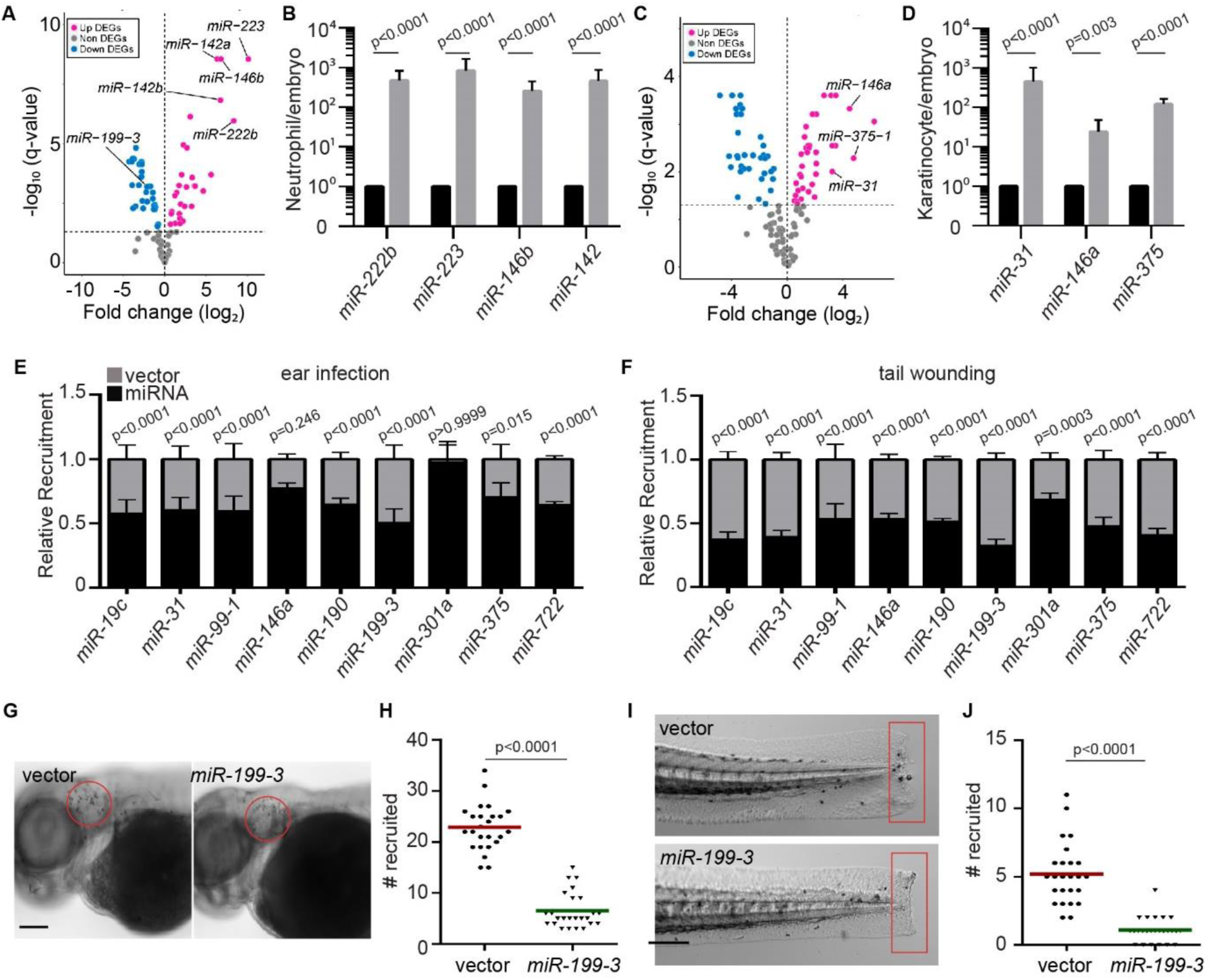
Identification of miRNAs that suppress neutrophil recruitment in vivo. (A) Differentially expressed genes (DEGs) of microRNAs in zebrafish neutrophils compared to whole embryo. (B) Quantification of selected miRs that were differentially expressed in neutrophils. (C) Differentially expressed genes (DEGs) of microRNAs in zebrafish epical keratinocytes compared to whole embryo. (D) Quantification of selected miRs that were differentially expressed in keratinocytes. Assay was done with 3 biological repeats each containing cells sorted from at least 100 larvae/repeat. Result is presented as mean ± SD, Holm-Sidak test. (E) Neutrophil recruitment to infected ear in transgenic lines with neutrophil specific overexpression of individual microRNAs. Results were normalized to number of neutrophils recruited in the vector expressing control lines in each individual experiment. (F) Neutrophil recruitment to tail fin transection sites in transgenic lines as described in c. (G, H) Representative images (G) and quantification (H) of neutrophils recruited to infected ear in vector or *miR-199-3* overexpressing zebrafish line. Scale bar, 100 µm. (I, J) Representative images (I) and quantification (J) of neutrophils recruited to tail fin transection sites in vector or *miR-199-3* overexpressing zebrafish line. Scale bar, 200 µm. (G-J) Assays were done with at least 2 individual founders with 3 biological repeats each containing 25 fish per group. Result is presented as mean ± s.d., Kruskal–Wallis test.

### miR-199-3 overexpression inhibits neutrophil motility and chemotaxis

To characterize the zebrafish lines *Tg(lyzC: Dendra2-miR-199-3*)*^pu19^* and *Tg(lyzC:Dendra2-vector)^pu7^*, hereby referred as *miR-199* and vector lines, we first isolated their neutrophils and performed miRNA RT-qPCR to confirm the *miR-199* overexpression in the *miR-199* line (fig. 2A). The levels of two abundant neutrophil miRNAs, *miR-223* and *let-7e*, were not affected, demonstrating that the overexpression did not disrupt physiological microRNA levels. Total neutrophil numbers in embryos were comparable between the two lines, suggesting that the observed decreased neutrophil recruitment was not an artifact of reduced neutrophil numbers (fig. 2B, C). In addition to defective recruitment to inflammatory sites, the velocity of neutrophil spontaneous motility in the head mesenchyme was significantly decreased in the *miR-199* line (fig 2D, E and Movie S1), whereas their directionality was not affected (fig. 2F). The sequences of the mature zebrafish and human *miR-199-5p* isoforms are identical (fig. 3C). We therefore sought to determine whether *MIR-199* inhibits human neutrophil migration. We used *MIR-100*, whose overexpression did not result in any phenotype in the initial screen, as a negative control. We expressed *MIR-199a*, *MIR-100* or the vector alone under the control of a Tet Responsive Element in HL-60 cells, a myeloid leukemia cell line that can be differentiated into neutrophil-like cells as our model (*24*) (fig. S2A). *MIR-100* and *MIR-199* were individually induced in the respective lines without affecting cell differentiation, cell death or the expression of two other miRNAs, *MIR-223* and *LET-7* (fig. 2G and fig. S2B-D). The cells were then allowed to migrate towards N-Formylmethionyl-leucyl-phenylalanine (fMLP) in transwells. Only the cells overexpressing *MIR-199*, but not the vector control or cells overexpressing *MIR-100*, displayed significant defects in chemotaxis (fig. 2H). Together, our results show that overexpression of *MIR-199* inhibits neutrophil migration in both zebrafish and human cell models.

**Fig. 2.**
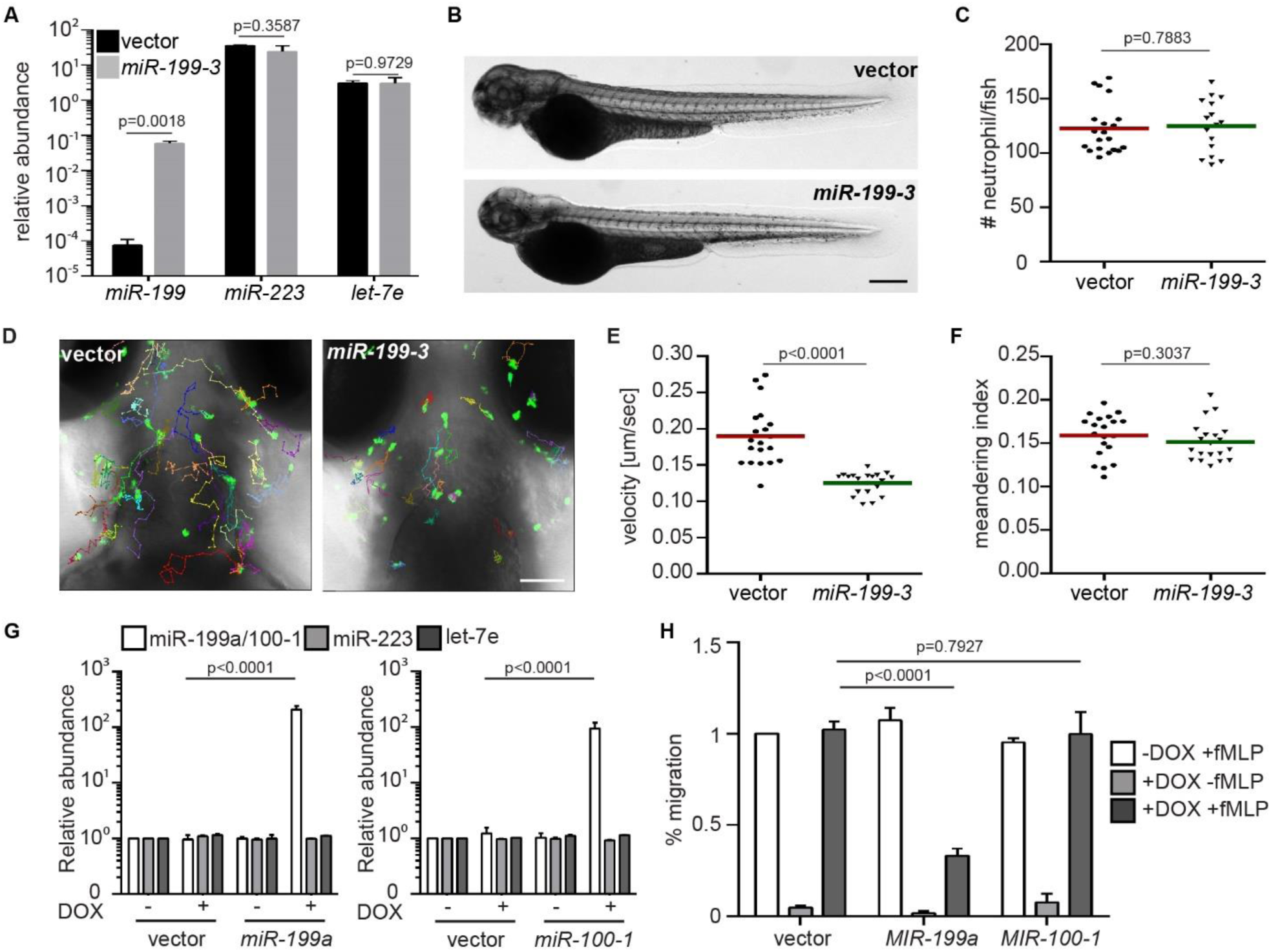
*miR-199-3* overexpression reduces neutrophil migration in zebrafish and human. (A) Quantification of *miR-199*, *miR-223* and *let-7e* levels in neutrophils sorted from the vector or *miR-199-3* zebrafish lines. Neutrophils were isolated from two adult kidney marrows from two different founders. Result are normalized to *u6* and presented as mean ± s.d., Holm-Sidak test. (B, C) Representative images (B) and quantification (C) of total neutrophils in vector or miR-199-3 zebrafish lines. Scale bar, 500 µm. Assays were done with 3 individual founders with 3 biological repeats each containing 20 fish per group. Result from one representative experiment is shown as mean ± s.d., Mann–Whitney test. (D-F) Representative images (D), velocity (E), and meandering index (F) of neutrophil motility in vector or *miR-199-3* zebrafish lines. Scale bar, 100 µm. 3 embryos each from three different founders were imaged and quantification of neutrophils in one representative video is shown. Kruskal–Wallis test. (G) Quantification of *MIR-100/199*, *MIR-233* and *LET-7E* in HL-60 cell lines with/without induced expression of the vector control or *MIR-100/199*. Result from three independent experiments are normalized to *U6* and presented as mean ± s.d., Holm-Sidak test. (H) Transwell migration of HL-60 cells with/without induced expression of the vector control, *MIR-199* or *MIR-100* toward fMLP. Results are presented as mean ± s.d., from three independent experiments and normalized to vector –DOX +fMLP, Kruskal–Wallis test.

**Fig. 3.**
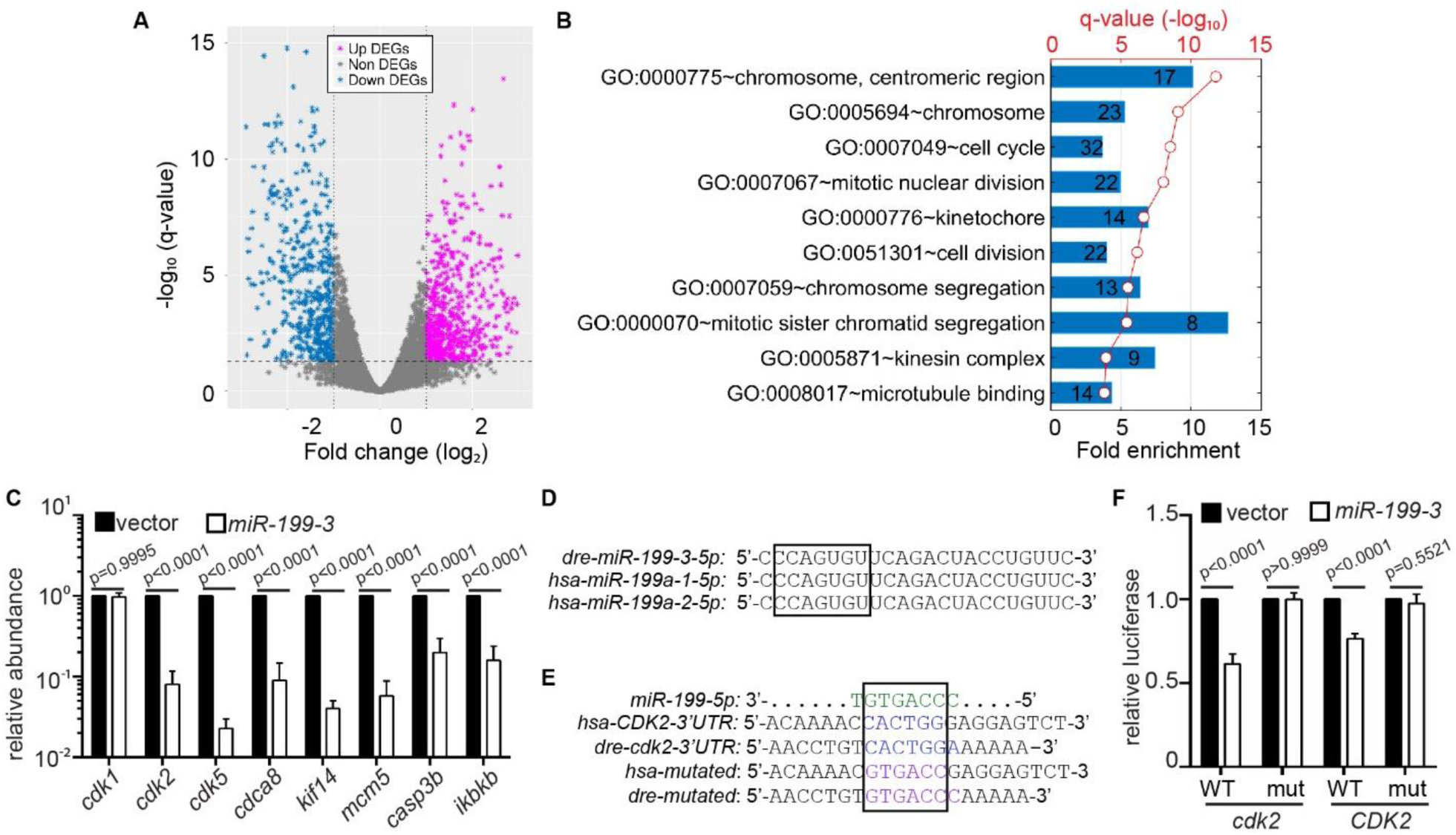
*miR-199-3* overexpression suppresses the expression of *Cdk2* in neutrophils. (A) Volcano blot of DEGs with significant expression changes in *miR-199* expressing neutrophils. Cyan: down-regulated differentially expressed genes (DEGs); Magenta: up-regulated DEGs. (B) Pathway analysis of genes downregulated by *miR-199* in neutrophils. (C) Quantification of the transcript levels of downregulated genes in neutrophils sorted from the vector or *miR-199-3* zebrafish line (at least 100 embryos per group/repeat). Results from three independent experiments are normalized to *rpl32* and presented as mean ± s.d., Holm-Sidak test. (D) Alignment of mature human and zebrafish *miR-199-5p* sequences. (E) Alignment of *miR-199-5p* and predicted *miR-199* binding sites in human and zebrafish *CDK2* 3’UTR. (F) Suppression of Renilla luciferase expressing via binding to both zebrafish and human CDK2 3’UTRs by *miR-199*. Results from three independent experiments are normalized to firefly luciferase and presented as mean ± s.d., Kruskal–Wallis test.

### miR-199 suppresses the expression of cell cycle-related genes including cdk2

To understand the underlying molecular mechanism of how *miR-199* regulates neutrophil migration, neutrophils from the *miR-199* or vector lines were isolated and their transcripts subjected to RNAseq. Transcripts which were downregulated by *miR-199* overexpression (*25*) were further analyzed (fig. 3A and Data file S4). We found that the top 10 pathways associated with the downregulated transcripts are related to cell cycle processes (fig. 3B and Data file S5). This result was surprising because neutrophils are known to be terminally differentiated, non-proliferative, and have exited the cell cycle (*26*). To validate the RNAseq results, we selected five genes, *cdk2, cdk5, cdca8, kif14*, and *mcm5*, which were significantly downregulated, and confirmed their reduced transcript levels in the *miR-199* line using RT-qPCR (fig. 3C). The zebrafish orthologue of *IKBKB*, a known target of human *MIR-199* (*27*) and a critical component in the NF-κB pathway, was also downregulated, supporting the conservation of microRNA-target interactions in zebrafish. The level of cyclin dependent kinase 1 (*cdk1*) mRNA was not altered. The protein sequence of CDK2 is more than 90% identical between zebrafish and human, and both 3’ UTRs contain predicted *miR-199-5p* binding sites (fig. 3D). To demonstrate direct targeting of *cdk2* mRNAs by *miR-199-5p*, we performed dual-luciferase reporter assays. *miR-199* significantly reduced the relative luciferase activity controlled by both the human and zebrafish *cdk2* 3’UTRs but did not show the repression of luciferase activity when the predicted binding sites were mutated (fig. 3E). These results indicate that *miR-199* suppresses classical cell cycle pathways in neutrophils and directly targets *cdk2* in both zebrafish and humans.

### Pharmacological inhibition of Cdk2 decreases neutrophil motility and chemotaxis

CDK2 is expressed and active in human neutrophils (*28*). To test whether *cdk2* regulates neutrophil migration, we used NU6140 (*29*), a CDK2 specific inhibitor. At a concentration of 100 µM, NU6140 did not affect zebrafish survival or neutrophil numbers (fig. S3). As a control for cell cycle progression and DNA replication, we used a mixture of Aphidicolin and hydoxyurea (ApH), as previously reported (*30, 31*) in an attempt to uncouple the role of Cdk2 from DNA replication. Neutrophil recruitment to the infected ear (fig. 4A, B) or tailfin amputation sites (fig. 4C, D) was significantly reduced by the Cdk2 inhibition, but not by DNA replication inhibition. Cdk2 inhibition did not affect total neutrophil numbers in the fish (fig. 5E, F). In addition, neutrophil motility, but not directionality, was significantly inhibited by the Cdk2 inhibitor (fig. 4G-I and Movie S2), recapitulating the phenotypes caused by *miR-199* overexpression. To increase our confidence of the results obtained with pharmacological Cdk2 inhibition, two additional Cdk2 inhibitors (*32*) were used. A CDK2 selective inhibitor CTV313 and a pan-CDK inhibitor roscovitine also blocked neutrophil motility and recruitment in inflammation models (fig. S4 and Movie S2). We then tested whether the neutrophils under investigation had indeed exited the cell cycle. No cell division was observed during wounding or infection (fig. S5A, B and Movie S3), suggesting that neutrophil division is a rare event. Additionally, less than 3% of the neutrophils in adult kidney marrow were in the S or G2 phase of the cell cycle (fig. S5C-E). Because *cdk2* has not been previously implicated in regulating neutrophil migration, primary human neutrophils were isolated and treated with NU6140. Inhibition of CDK2 significantly reduced chemotaxis toward Interleukin-8 (IL-8), with a decreased velocity and chemotaxis index (fig. 4J-L and Movie S4). In summary, we report a previously unknown role for CDK2 in regulating neutrophil chemotaxis in a way separable from its function in DNA replication.

**Fig. 4.**
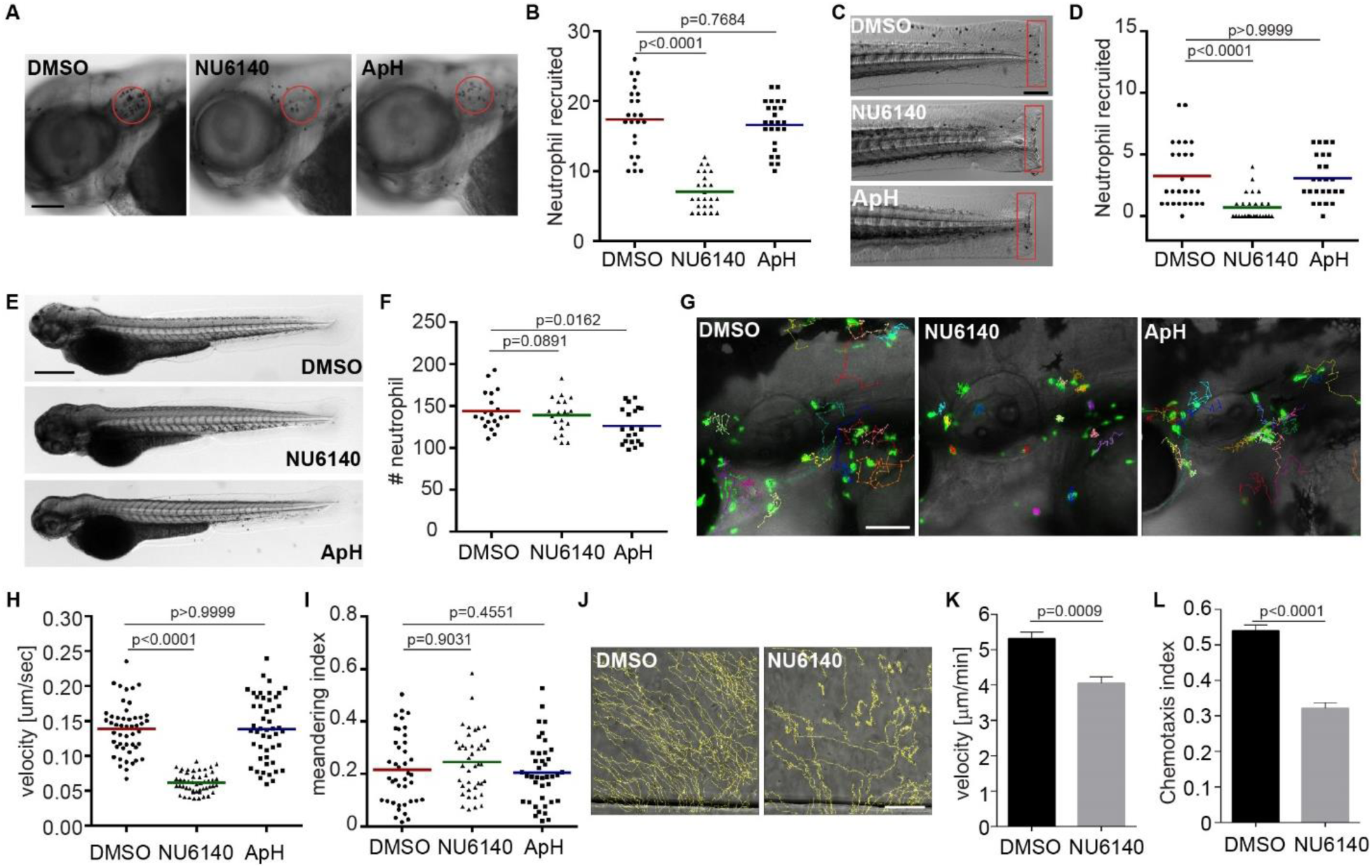
Inhibition of CDK2 reduces neutrophil motility and chemotaxis in zebrafish and humans. (A, B) Representative images (A) and quantification (B) of neutrophils recruited to infected ear in zebrafish larva treated with cdk2 inhibitor (NU6140) or DNA replication inhibitors Aphidicolin +hydoxyurea (ApH). Scale bar, 100 µm. (C, D) Representative images (C) and quantification (D) of neutrophils recruited to tail fin transection sites in zebrafish larva treated with NU6140 or ApH. Scale bar, 200 µm. (E, F) Representative images (E) and quantification (F) of total neutrophil number zebrafish larva treated with NU6140 or ApH. Scale bar: 500 µm. (A-F) Assays were done with 3 individual founders with 3 biological repeats each containing 20 (for motility) or 25 (for neutrophil recruitment) fish per group. Result from one representative experiment is shown as mean ± s.d., Mann–Whitney test. (G-I) Representative images (G), velocity (H), and meandering index (I) of neutrophil motility in zebrafish larvae treated with NU6140 or ApH. Scale bar, 100 µm. 3 embryos each from three different founders were imaged and quantification of neutrophils in one representative video is shown, Kruskal–Wallis test. (J-L) Representative tracks (J), mean velocity (K), and chemotaxis index (L) of primary human neutrophils treated with DMSO or NU6140 (50 µM) migrating towards IL-8. Scale bar, 100 µm. Results representative for 3 separate trials are shown. Result is presented as mean ± s.e.m., Two-way paired Welch’s t-test.

**Fig. 5.**
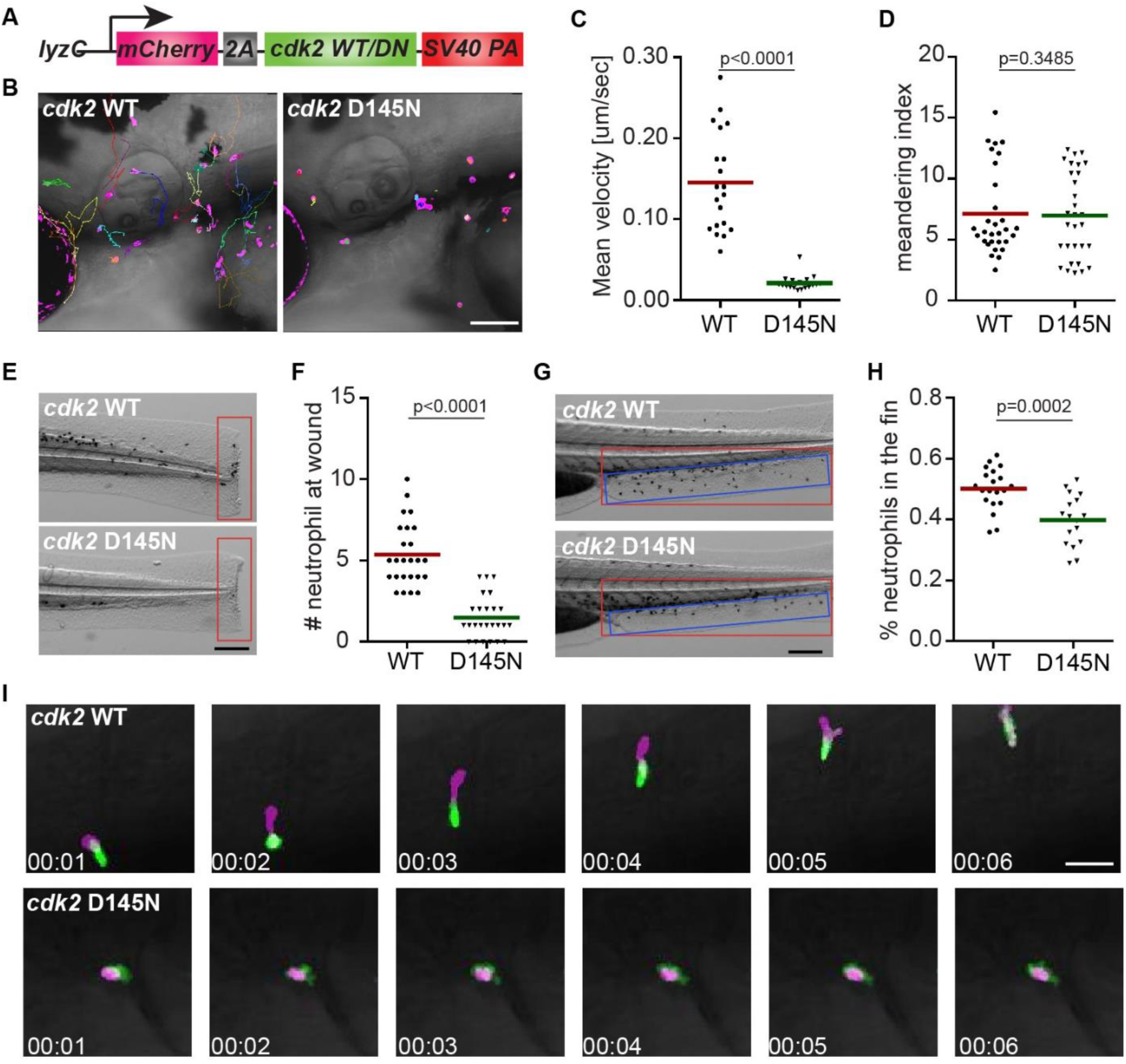
Dominant-negative CDK2 suppresses neutrophil motility and chemotaxis. (A) Schematic of the construct for neutrophil specific expression of CDK2 wild-type (WT) or D145N. (B-D) Representative images (B), velocity (C), and meandering index (D) of neutrophil motility in CDK2 WT or D145N zebrafish line. Scale bar, 100 µm. 3 embryos each from three different founders were imaged and quantification of neutrophils in one representative video is shown, Kruskal–Wallis test. (E, F) Representative images (E) and quantification (F) of neutrophil recruitment to the tailfin transection sites in CDK2 WT or D145N zebrafish line. Scale bar, 200 µm. (G, H) Representative images (G) and quantification (H) of neutrophils migrated to the caudal fin (blue box) normalized to total neutrophils in the trunk (red box) in CDK2 WT or D145N zebrafish line. Scale bar, 200 µm. (I) Simultaneous imaging of utrophin-GFP distribution in neutrophils expressing either WT or D145N Cdk2. Data are representative of more than three separate time-lapse videos. Scale Bar, 50µm. (E-H) Assays were done with 3 individual founders with 3 biological repeats each containing 20 (for motility) or 25 (for neutrophil recruitment) fish per group. Result is presented as mean ± s.d., Mann–Whitney test.

### Catalytic activity of Cdk2 is required for neutrophil motility

We further tested our hypothesis that the kinase activity of Cdk2 is essential for neutrophil motility. We cloned the zebrafish *cdk2* gene and tagged it at the N-terminus with mCherry linked via a self-cleavable 2a peptide (fig. 5A). Mutations at D145, a residue that chelates Mg^2+^ and facilitates ATP binding to the catalytic cleft (*33*), resulted in a catalytic-dead and dominant-negative form of Cdk2. We constructed zebrafish lines overexpressing *cdk2* (D145N) in neutrophils, *Tg(lyzC:mcherry-2a-cdk2-d145n)^pu19^*, together with a control line *Tg(lyzC:mcherry-2a-cdk2)^pu20^*, hereby referred to as cdk2 D145N or the WT line. We observed a significant reduction of neutrophil motility (fig. 5b-D and Movie S5) and recruitment to the tail amputation site (fig. 5E, F) in the D145N line. To characterize neutrophil chemotaxis to a single chemoattractant, we bathed embryos from the two lines in Leukotriene B4 (LTB4) and quantified neutrophil emigration from the caudal hematopoietic tissue (CHT) to the caudal fin. The percentage of neutrophils migrating outside the CHT and the distance neutrophils traveled in the caudal fin were significantly reduced in the D145N line (fig. 5G, H, fig. S6A, B and Movie S6). In addition, neutrophil motility in the CHT was reduced (fig. S6C, D and Movie S7). To visualize neutrophil cell polarity, the cdk2 D145N or the WT lines were crossed with *Tg(mpx:GFP-UtrCH*) which labels stable actin in the trailing edge of migrating neutrophils (*34, 35*). In Cdk2 D145N expressing neutrophils, stable actin is enriched in transient cell protrusions that quickly retract in multiple directions, displaying disrupted cell polarity and actin dynamics (fig.5I and Movie S8). Together, our results indicate that the catalytic activity of Cdk2 is required for neutrophil polarization and chemotaxis.

### miR-199 overexpression or Cdk2 inhibition ameliorates systemic inflammation

With the findings that *miR-199* overexpression or *cdk2* abrogation hinders neutrophil chemotaxis, we sought to determine whether these manipulations could alleviate acute systemic inflammation caused by *P. aeruginosa* infection (*36, 37*). We thus administered a lethal dose of *P. aeruginosa* directly into the circulation in zebrafish embryos. Survival of the embryos was significantly improved in the *miR-199* line and the *cdk2* D145N line compared with their respective control lines (fig. 6A, B). Consistently, NU6140 treatment improved the infection outcome whereas DNA replication inhibitors were detrimental to the infected host (fig. 6C). Roscovitine, a pan CDK inhibitor (*38*) currently in phase-II trial for cancer treatment, also displayed a protective effect at 15 µM but was detrimental at 30 µM, possibly due to the inhibitory effect on cell proliferation and tissue repair as previously reported in cell culture and *ex vivo* bone marrow isolations (*39, 40*) (fig. 6D). To assess further the protective effect of dampening neutrophilic inflammation during systemic inflammation, we utilized a sterile inflammation model of endotoxemia (*41*). Neutrophil-specific *miR-199* overexpression or roscovitine treatment reduced host mortality (fig. 6E, F), indicating a host protective effect of *miR-199* overexpression and Cdk2 inhibition during acute systemic inflammation and infection, without generating broad immune suppression.

**Fig. 6.**
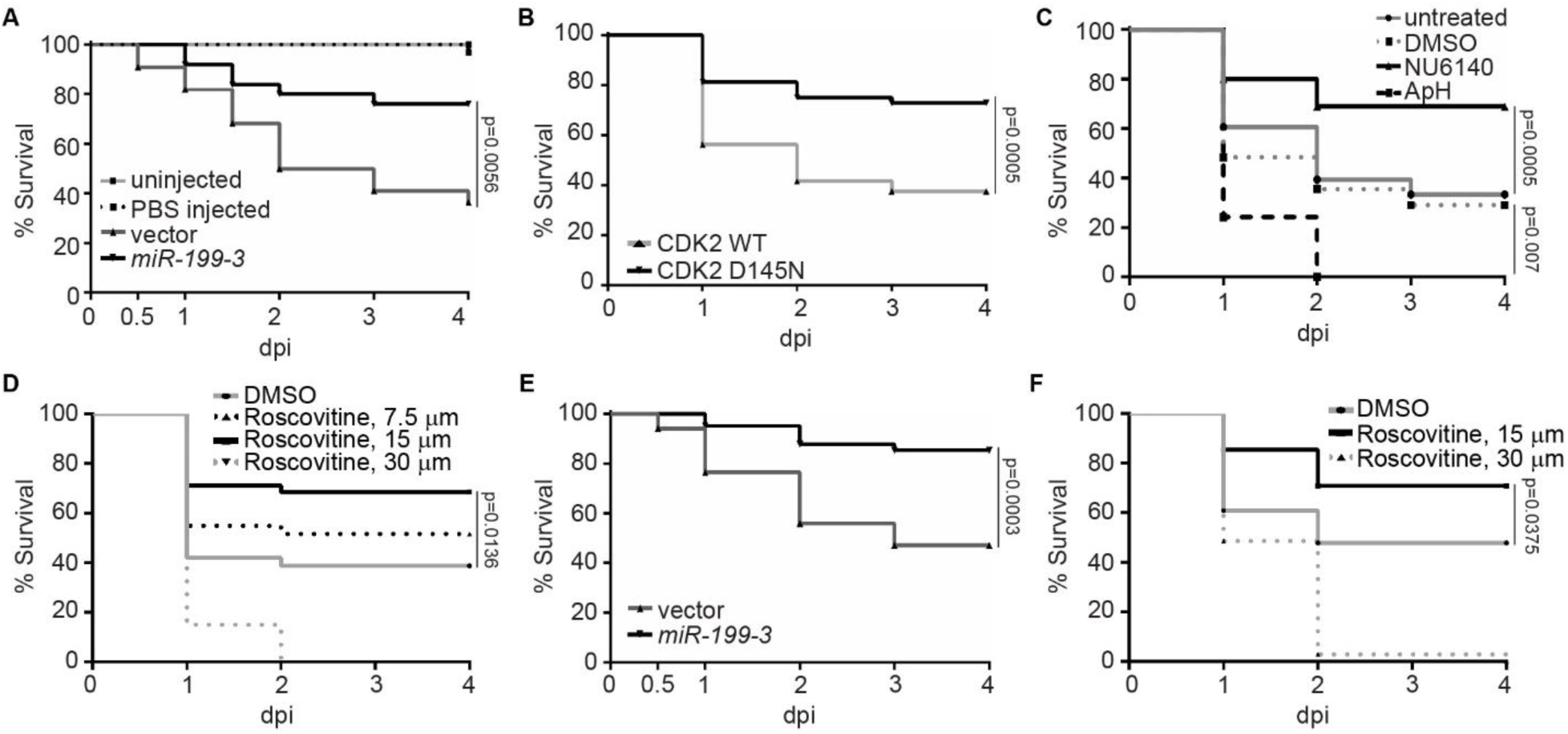
Cdk2 inhibition and *miR-199* overexpression improves zebrafish survival during systemic bacterial infection or sterile inflammation. (A) Survival of zebrafish larvae after infection of 1000 CFU of *P. aeruginosa* i.v in the vector or *miR-199* zebrafish line. Vector lines were left uninfected or injected with PBS as a control. (B) Survival of zebrafish larvae after infection of 1000 CFU of *P. aeruginosa* i.v in the CDK2 WT or D145N zebrafish line. (C) Survival of zebrafish larvae treated with vehicle, NU6140 (100 µM), ApH (150 µM/20 mM) or left untreated after infection of 1000 CFU of *P. aeruginosa* i.v. (D) Survival of zebrafish larvae treated with indicated dilutions of roscovitine infected with 1000 CFU of *P. aeruginosa* i.v.. (E) Survival of zebrafish larvae after injection of 25 ng LPS i.v. in the vector or *miR-199* zebrafish line. (F) Survival of zebrafish larvae treated with indicated dilutions of roscovitine injected with 25ng LPS i.v.. (A-F) One representative experiment of three independent experiments (n >=20 each group) is shown, Gehan–Breslow–Wilcoxon test.

## Discussion

microRNAs play a pivotal role in diseases driven by neutrophilic inflammation (*42–44*) where both protective and detrimental effects are observed. Currently, there is no systemic screen to identify miRNAs that suppress neutrophil migration, possibly because neutrophils are terminally differentiated and not genetically tractable in cell culture. A microRNA overexpression screen was performed in B cells (*15*), but not in other immune cells. In the present study, we performed an *in vivo* functional screen in the zebrafish and identified *miR-199* as a critical evolutionarily conserved regulator of neutrophil migration and chemotaxis. *miR-199* hindered neutrophil motility by downregulating genes in the canonical cell cycle pathways, and by directly targeting Cdk2. We further exhibited that the catalytic activity of CDK2 is critical for neutrophil migration and chemotaxis, whereas DNA replication and an active cell cycle are not essential (fig. S6E).

Whereas no previous studies have investigated the role of *miR-199* in neutrophils, its role in suppressing inflammation and cell migration has been reported in cancer cells. In ovarian cancer cells, *miR-199* inhibits canonical NF-κB signaling by targeting *ikbkb*, thus preventing the production of pro-inflammatory cytokines (*27*). In our current work, the *miR-199*-*ikbkb* axis was also validated in neutrophils, further supporting the evolutionary conservation of miRNA-target interaction in vertebrates. Suppression of *ikbkb* may also contribute to the survival of zebrafish during systemic inflammatory challenge due to its central role in inflammation. From the perspective of cell migration, *miR-199* hinders cancer cell migration and invasion by downregulating β1 integrin levels (*45*), decreasing CD44 and interaction with Ezrin (*46*), or by directly targeting α3 integrin (*47*) in various cancer cells. Low levels of *miR-199* correlate with poor prognosis, increased cell proliferation and metastasis. Subsequent introduction of *miR-199* alleviated invasiveness *in vitro* through targeting the Wnt or HIF-1α/VEGF pathways (*48, 49*). Here we provide evidence that *miR-199* is a suppressor of cell migration in leukocytes, expanding its role beyond inflammation. Our observation is in line with recent reports that *miR-199a-5p* is increased in the plasma in patients with neutrophilic asthma (*50*) and inflammatory bowel disease (*51*), suggesting that miR-199 may play physiological functions in inflammation. In addition, *miR-199b* suppress myeloid differentiation (*52*), which can also contribute to its role in suppressing the innate immunity.

The most surprising finding of our study is that *miR-199* predominantly regulates the cell cycle-dependent kinase *CDK2* in terminally differentiated neutrophils that have exited the cell cycle. The function of CDK2 has not previously been investigated in the context of inflammation. CDK2 is a cell cycle-dependent kinase whose cofactors, cyclin E and A, are expressed during mitosis, activating its kinase activity and cell cycle progression (*53*). CDK2 is a facilitator of the cell cycle during G1/S phase (with cyclin E) and S/G2 phase (with cyclin A) (*54*), and deletion of CDK2 prolongs but does not inhibit cell cycle progression (*55*). CDK2 is required for meiosis during mammalian gametogenesis. In addition, CDK2 phosphorylates transcriptional factors to drive cell cycle progression and cell differentiation, and to alter cell metabolism and DNA double-strand break repair (*56*).

We provided evidence that the kinase activity of CDK2 is required for neutrophil migration. The limitation of our work is that the direct phosphorylated substrate of CDK2 in neutrophils are not identified, which presents a significant challenge. CDK2 is a serine/threonine protein kinase, which uses the KLAD*FGLA kinase consensus domain to phosphorylate the “Ser/Thr-Pro-X-basicAA” motif of target proteins, although noncanonical target motifs are prevalent (*57*). Over a hundred proteins are likely phosphorylated by CDK2-cyclinA in HEK cells (*58*) and the CDK2-cyclinE substrates are not well characterized. Thus extensive work will be required to characterize how CDK2 contributes to the signaling network required for neutrophil chemotaxis and identify the proteins that are phosphorylated by CDK2 in neutrophils. CDK2 is present in the neutrophil cytosol and its expression is upregulated upon LPS stimulation (*59*), supporting the possibility that CDK2 ties in the signaling pathways downstream of the immune receptors in neutrophils. As a step closer to the potential mechanism, we have visualized the actin dynamics and the cell polarity in neutrophils with the dominant-negative Cdk2 overexpression. Indeed, stable actin is enriched in the transient protrusions that quickly retract, a phenotype resembling the over expression of a dominant negative Rac2 construct and after PI3K inhibition (*35, 60*). It is likely that Cdk2 regulates the Rac2-PI3K axis, which is essential in cell motility.

Acute neutrophilic over-inflammation is a pressing concern in many disease pathologies (*1*) and methods that simultaneously hinder neutrophil migration and inflammation while preserving immune integrity are highly desired. Our study presents an acute method to dampen neutrophil recruitment by inhibiting Cdk2, which reduces the lethality caused by acute systemic inflammation. The pan-CDK inhibitor roscovitine competes with ATP for the occupation of the kinase active site (*61*) and is currently in multiple phase II clinical trials to inhibit cystic fibrosis and multiple types of tumors (clinicaltrials.gov). Previous studies in mice have shown that treatment of roscovitine at 10-100 mg/kg for 3-7 days after inflammation induction resulted in decreased neutrophil infiltration into the inflamed tissue and better clinical outcome (*62*). This reduced neutrophil recruitment was attributed to CDK9 inhibition and neutrophil apoptosis (*63*) and a direct link to CDK2 was not shown. *Cdk2* knockout mice show normal myeloid cell numbers during resting and stressed conditions (*64*) and no developmental defects other than having a smaller body size and sterility (*65*). Thus CDK2 inhibition is not expected to cause adverse side effect and supports an attractive strategy for short-term and long-term control of inflammation. Therefore, our results here provide a mechanistic understanding of the neutrophil inhibitory effect of abrogating CDK2 and suggest a future direction for utilizing Cdk2 specific inhibitors in managing inflammation, which may be better tolerated than a pan-Cdk inhibitor such as roscovitine. In summary, our present study demonstrates previously unappreciated roles of *miR-199* and CDK2 in regulating neutrophil migration and points to new directions in managing neutrophilic inflammation.

## Materials and Methods

### Study Design

The objective of this research is to identify new molecules that regulate neutrophil migration. The zebrafish experiment was conducted in accordance with internationally accepted standards. The Animal Care and Use Protocol was approved by The Purdue Animal Care and Use Committee (PACUC), adhering to the Guidelines for Use of Zebrafish in the NIH Intramural Research Program (protocol number: 1401001018). We used MATLAB and the samplesizepwr function to calculate the sample sizes required for each experiment based on conservative estimates for the variability in the controls for each type of experiments. With a power of 0.9 (significance level of 0.05) in two sample t test, we need a sample size of 15 neutrophils to detect a change of 20% in the motility assay; 11 embryos to detect a change of 20% in LTB4 fin recruitment assay; 11 embryos to detect a change of 20% in total neutrophil numbers; 22 and 25 embryos to detect a 50% change of neutrophil numbers recruited to the wound and ear infection sites respectively. Results were derived from multiple founders for each transgenic line. All data were included in final analysis and quantified blindly. All experiments were repeated at least three times.

## Authorship Contributions

AYH, AH, DU, JW and DQ designed research and wrote the manuscript. AYH, DW, JL, DB performed the experiments. AYH, DW, JL, JW, RS, DB, DU and SL analyzed data. All authors read and approved the manuscript.

## Data and materials availability

The microRNA-seq and RNA-seq raw and processed data are submitted to the Gene Expression Omnibus (GSE127174). Plasmids are available on Addgene.

## Supplementary Materials

Fig. S1. miRNA screen experimental design.

Fig. S2. Characterization of dHL-60 cell lines with inducible miR overexpression.

Fig. S3. Cytotoxicity of the CDK2 inhibitor NU6140.

Fig. S4. The CDK2 inhibitor CTV313 or a pan-CDK inhibitor Roscovitine reduces neutrophil migration.

Fig. S5. Cell cycle profiling of zebrafish neutrophils.

Fig. S6. Cdk2 D145N inhibits neutrophil motility and chemotaxis and the working model.

Data File S1. Relative expression of microRNAs in sorted zebrafish neutrophils and apical keratinocytes compared to whole embryos at 3 dpf.

Data File S2. Quantification of percentage of neutrophils recruited to the infected ear from the head in upon transient expression of selected miRNAs.

Data File S3. Quantification of number of neutrophils recruited to the infected ear and tail wounding in stable lines.

Data File S4. List of genes that are downregulated in the *miR-199* overexpressing neutrophils.

Data File S5. Pathway enrichment analysis of downregulated genes in *miR-199* overexpressing neutrophils.

Movie S1: Tracked movies of neutrophil motility from the vector and the *miR-199* lines in the head mesenchyme.

Movie S2: Tracked movie of neutrophil motility in the head mesenchyme treated with vehicle, NU6140, Aphidicolin+hydoxyurea, CTV313, or Roscovitine.

Movie S3: Neutrophils are not dividing when recruited to regional inflammation sites.

Movie S4: NU6140 inhibits chemotaxis of Primary human neutrophil.

Movie S5: Tracked movies of neutrophil motility from the Cdk2 WT and D145N lines in the head mesenchyme.

Movie S6: Tracked movies of neutrophil recruitment to the fin in the Cdk2 WT and D145N lines bathed in LTB4.

Movie S7: Tracked movies of neutrophil motility in the Cdk2 WT and D145N lines in the caudal hematopoietic tissue.

Movie S8: Morphology of neutrophils from the Cdk2 WT and D145N lines.

