## supplemental methods and figures for "Phenotypical microRNA screen reveals a noncanonical role of CDK2 in regulating neutrophil migration"

### Supplementary materials and methods

*microRNA seq*

Neutrophil and apical epithelial cells were sorted from transgenic zebrafish lines Tg(lyzC:GFP) and Tg(Krt4:GFP) at 3 day post fertilization (dpf) using fluorescence-activated cell sorting. Total RNA was exacted using Ambion mirVana kit (without enrichment for small RNA). Library was prepared using TruSeq Small RNA Preparation (illumina). After library construction, DNA of 145-160 bp were excited from Invitrogen 6% Novex TBE Gel and sequenced using Illumina HiSeq2000 with paired-end reads of 50 bp with a total of 2-3M/sample. MicroRNA sequence is performed at University of Wisconsin Collaborative Genomics Core. For analysis, sequencing reads were map to the zebrafish genome (GRCz11) using RNA-seq aligner from STAR (v2.5) (66) with the parameter: "--alignEndsType EndToEnd --outFilterMismatchNmax 1 --outFilterMultimapScoreRange 0 --outFilterMultimapNmax 10 --outSAMunmapped Within --outFilterScoreMinOverLread 0 --outFilterMatchNminOverLread 0 --outFilterMatchNmin 16 --alignSJDBoverhangMin 1000". FeatureCounts (from subread v1.5.1) *(67)* was used to evaluate expression levels of zebrafish mature microRNAs based on uniquely mapped reads with the parameters: "-s 1 –Q 10". Before TMM normalization and DE analysis by edgeR (v3.20.8) *(68)*, microRNAs whose CPM was less than 10 were removed from all samples. Differentially expressed microRNAs were selected if their FDR-adjusted p-values were less than 0.05.

*RNA seq*

Kidney marrow was dissected from 2 adults from Tg(lyzC:Dendra2-miR-199-3)pu19 or Tg(lyzC:Dendra2-vector)pu7 and neutrophils were sorted using fluorescence-activated cell sorting. Total RNA was extracted using RNeasy Plus Mini Kit (Qiagen #74104). RNAseq was performed at The Center for Medical Genomics at Indiana University School of Medicine. Samples were polyA enriched and sequenced with Illumina HiSeq 4000 ultra low with reads range from 37M to 44M. The RNA-seq aligner from the STAR (v2.5) *(66)* were employed to map RNA-seq reads to the reference genome, zebrafish (GRCz11), with the following parameter: "--outSAMmapqUnique 60". Uniquely mapped sequencing reads were assigned to genes using featureCounts (from subread v1.5.1) *(69)* with the following parameters: "-p -Q 10". The genes were filtered for further analysis if their count per million (CPM) of reads was less than 0.5 in more than 3 samples. The method of trimmed mean of M values (TMM) was adopted for gene expression normalization cross all samples, followed by differential expression analysis between different conditions using edgeR (v3.20.8). Differentially expressed gene was determined for the comparison if its false discovery rate (FDR) adjusted p-value was less than 0.05 and the amplitude of fold change (FC) was larger than linear 2-fold. The functional analysis was performed on DEGs of our interest with a cutoff of FDR < 0.05 to identify significantly over-represented Gene Ontology (GO) and/or KEGG pathways by using the DAVID *(70).*

*Quantitative RT-PCR:*

Total RNA was purified using MiRVANA miRNA purification kit (ThermoFisher). MicroRNAs were reverse transcribed with Universal cDNA Synthesis Kit II (Exiqon). MicroRNA RT-qPCR was performed with ExiLENT SYBR® Green master mix (Exiqon) using LightCycler ® 96 Real-Time PCR System (Roche Life Science). Messenger RNAs were reverse transcribed with Transcriptor First Strand cDNA Synthesis Kit (Roche). RT-qPCR were performed with FastStart Essential DNA Green Master (Roche). The specificity of the primers were verified with a single peak in the melt-curve. The relative fold change with correction of the primer efficiencies was calculated following instructions provided by Real-time PCR Miner (http://ewindup.info/miner/data_submit.htm) and normalized to U6. miRNA Primers used in this study are: dre-miR-222b (2111861), dre-miR-146b (2117034), dre-miR-142 (2104518), dre-miR-146a (2114679), dre-miR-21 (2101053), dre-miR-430 (2105859), dre-miR-92a (204258), dre-miR-181a (2100735), dre-miR-31 (2103532), dre-miR-375 (2109706), dre-miR-199-3 (204536), dre-miR-223 (205986), dre-let7e (2106780), dre-U6 (206999), hsa-miR-199 (204536), hsa-miR-100 (205689), hsa-miR-223 (205986), hsa-let7e (205711), hsa-U6 (203907).For One-step RT-qPCR of sorted neutrophils, RNA was extracted as using RNeasy Plus Mini Kit (Qiagen #74104) and one-step RT-qPCR was performed with SuperScript® III Platinum® SYBR® Green One-Step qRT-PCR Kit (Invitrogen), using LightCycler® 96 Real-Time PCR System (Roche Life Science). The relative fold change with correction of the primer efficiencies was calculated following instructions provided by Real-time PCR Miner and normalized to rpl32. Primers: dre-cdk1+: 5'-ttggggtcccagtaagagtcta-3'; dre-cdk1-: 5'-ggtgtggaatagcgtgaagc-3'; dre-cdk2+: 5'-agatggcagagaccttctcg-3'; dre-cdk2-: 5'-cggaaaaaccgatgaacaag-3' ; dre-cdk5+: 5'-tgggaacgccaacagaag-3'; dre-cdk5-: 5'- agctggatacatcgggtacg-3' ; dre-cdca8+: 5'-atggcaccgctgaagtctac-3'; dre-cdca8-: 5'-cgttgaaggagctttctttga-3' ; dre-kif14+: 5'-aggaggctcgcttgaagg-3'; dre-kif14-: 5'-ctctttagccacctgaatgc-3' ; dre-mcm5+: 5'-aagctattgcctgcctgct-3'; dre-mcm5-: 5'-ccctcttcgtgtcagaccat-3' ; dre-casp3b+: 5'-caggatattacgcatggagga-3'; dre-casp3b-: 5'- tcgcacagcgaggagataa-3' ; dre-ikbkb+: 5'-gaggagcgagaaacaactgc-3'; dre-ikbkb-: 5'-tggactgcgaactttactgc-3' ; dre-rpl32+: 5'-tcagtctgaccgctatgtcaa-3'; dre-rpl32-: 5'- tgcgcactctgttgtcaatac-3'.

*Generation of transgenic zebrafish lines*

300-600bp genomic DNA fragments flanking the respective miRNA were PCR amplified from zebrafish genomic DNA using specific primers and inserted into the BbsI site in the vector backbone as described (22). Zebrafish cdk2 gene was cloned from zebrafish mRNA using SuperScript III RT-kit (Invitrogen #18080044) and amplified with the following primers. zCdk2+: 5´- GAAAACCCCGGTCCTATGGAGTCTTTTCAGAAAGTGGAG-3´; zCdk2-: 5´- CATGGCTGATTATGATTATAGGCGTAAAGGAGGCACTGG-3´ and inserted into a Tol2-mcherry-2A backbone. Point mutation was generated using infusion (Takara #638920) site directed mutagenesis with zCdk2 DN+: 5'- ACTGGCTAACTTTGGTTTGGCCAGAGCGTTC -3'; zCdk2 DN -: 5'- CCAAAGTTAGCCAGTTTGATCTCGCCCTG -3'. The plasmids were injected into zebrafish embryo at 1-cell stage and the stable line were generated as previously described (35). At least two founders (F0) for each line were obtained. Experiments were performed using F2 larvae produced by F1 fish derived from multiple founder to minimize the artifacts associated with random insertion sites.

Dual luciferase reporter assay:

Both zebrafish and human CDK2 3´UTR were amplified from zebrafish genomic DNA with SuperScript III RT-kit (Invitrogen #18080044) and cloned into psiCHECK2 (Promega) at XhoI and NotI cloning sites with the following primers: zCDK2+: 5'-TAGGCGATCGCTCGAGAACGAGATCAACTTTGGCAAG-3'; zCdk2-: 5'-TTGCGGCCAGCGGCCGCTGTAACAACATAAACCAAATGTT-3', hCDK2+: 5'-TAGGCGATCGCTCGAGAGCCTTCTTGAAGCCCCCA-3'; hCdk2-: 5'-TTGCGGCCAGCGGCCGCTATAAAACTAGGCACATTTTTTTAA-3'. Mutated 3'UTR constructs were generated using Infusion HD cloning kit (Clontech) with the following primers: zCdk2 mut+: 5'- CCTGTGTGACCCAAAAAAACAGCTGTGCTAATAGT-3'; zCdk2 mut-: 5'- TTTGGGTCACACAGGTTGCAATGGATAATCTTCA-3', hCdk2 mut+: 5'-AAAACGTGACCGAGGAGTCTATTTTAAAGAATTCGG. DNA encoding miR-199 or vector were amplified from the construct used for expression in zebrafish and inserted into pcDNA3.1 at the HindIII/XbaI cloning sites using the following primers: pcDNA-199+: 5'-GTTTAAACTTAAGCTTGCCACCATGGATGAGGAAATCGC-3'; pcDNA-199-: 5'-AAACGGGCCCTCTAGAGACCGGTACCCCCGGGCTGC-3'.

*Live imaging:*

Larvae at 3 dpf were settled on a glass-bottom dish and imaging was performed at 28.5 ºC. Time-lapse fluorescence images in the head mesenchyme were acquired with a laser-scanning confocal microscope (LSM710, Zeiss) with a Plan-Apochromat 20×/0.8 M27 objective. Neutrophil motility at the caudal hematopoietic tissue was imaged using Zeiss EC Plan-NEOFLUAR 10X/0.3 objective. For neutrophil nucleus and cytosol reporter line imaging, a LD C-Apochromat 40x/1.1 W Korr M27 was used. The green and red channels were acquired sequentially with 0.1~0.3% power of the 488 nm laser and 0.5%~2% of 561 nm laser, respectively, with a 200 µm pinhole at a speed of 1.27 µs/pixel and averaged (line 2). The fluorescent stacks were flattened using the maximum intensity projection and overlaid with a single slice of the bright-field image. Neutrophil chemotaxis upon LTB4 treatment was captured with a Zeiss ZV16 dissection microscope at 2x magnification every 15sec for 30 minutes.

*Inflammation assays in zebrafish:*

Zebrafish wounding and infection were performed as described *(22).* For LTB4 recruitment assays, 3 dpf zebrafish larvae were treated with 30 nM LTB4 for 15 minutes and fixed. Neutrophils were stained with Sudan Black and the number at the indicated regions were quantified.

*Generation of stable HL-60 cell lines:*

HEK-293 cells were cultured in DMEM supplemented with 10% FBS, 4.5 g/glucose and sodium bicarbonate. HL-60 cells were obtained from ATCC (CCL-240) and cultured using RPMI-1640 with HEPES supplemented with 10% FBS and sodium bicarbonate. The lentiviral backbone pLIX_403 was a gift from David Root (Addgene plasmid # 41395). The DNA sequence flanking MIR-199 was cloned from HL-60 cell genomic DNA and cloned into a backbone containing

Dendra2. The miRNA and dendra2 reporter were then cloned into pLIX_403 vector using the NheI/AgeI sites with primer set: pLIX-mir+: 5' –TGGAGAATTGGCTAGCGCCACCATGG ATGAGGAAATCGC-3', pLIX-mir-: 5'-CATACGGATAACCGGTTACCACACCTGGCTGG GC-3'. Stable HL-60 cell lines were generated as described *(71).* Briefly, HEK-293 cells were transfected with pLIX_403, VSV-G and CMV8.2 using lipofectamine 3000 (ThermoFisher). The viral supernatant was harvested on day three and concentrated with Lenti-X concentrator (Clontech). HL-60 Cells were infected at 2000 g for 2 h at 32 ºC and maintained in medium supplemented with 1 µg/ml puromycin. Live cells that excluded trypan blue were counted using a hemocytometer.

*Flow Cytometry:*

Cells were stained with Alexa Fluor 647 conjugated CD11b (neutrophil differentiation marker, Biolegend, 301319, clone IV-M047, 2ul to 1X106 cells) or the isotype control antibody (Biolegend, 400130, clone MOPC-21, 2ul to 1X106 cells) and RUO AnnexinV (apoptosis marker, BD, 563973, 5ul to 1X106 cells). Cells were stained in staining buffer (1% BSA, 0.1% NaN3) and incubated on ice for 1 hour, washed three times with staining buffer and resuspended in suitable volumes. Cell cycle profiling was obtained using Vybrant™ DyeCycle™ Ruby Stain (Thermo #V10309) following the manufacture's protocol. Fluorescence intensity were collected using BD LSR Fortessa. Results were analyzed with Beckman Kaluza 2.1 software.

*Inhibitor treatment for zebrafish larvae:*

CDK2 inhibitors NU6140 (Enzo # ALX-270-441), CTV313 (Sigma #238803) and roscovitine (Selleckchem #S1153) were dissolved in DMSO to make a 100 mM stock, then further diluted in E3 to working concentrations (100 µM for recruitment and motility assays, and indicated concentration for survival assays). For inhibiting DNA replication, 150 µM aphidicolin and 20 mM hydroxyurea (Sigma #A0781) in 1% DMSO was used as previously described *(30, 72).* For neutrophil recruitment and random motility assays, larval were pretreated with the inhibitor for 1 h before experimental procedures. For survival assays, larval were pretreated and kept in the inhibitors from1 hour post infection.

*Primary neutrophil isolation and chemotaxis assay:*

Primary human neutrophils were isolated with Milteny MACSxpress Neutrophil Isolation Kit. Cells were stain for 10 min with Calcein AM and washed with PBS. 10 million cells/ml were incubated in RPMI+0.1% HSA containing 50 µM NU6104 or DMSO at 37ºC/5% CO2 for 45 min. Microfluidic devices were fabricated as previously described *(73).* Microfluidic chambers were coated with 10 µg/ml fibrinogen for 1 h at 37 °C in PBS. 3 µl of neutrophil suspension (4 × 106 cells/ml) was added to the device, and 5 µM IL-8 was added to the source port. The gradient was allowed to set up and equilibrate in a 37 °C humidified chamber for 15 min before imaging. Time-lapse imaging was performed using a 10×, NA of 0.45, or 60×, NA of 1.4 objective and motorized stage (Ludl Electronic Products) on an inverted microscope (Eclipse TE300) using a charge-coupled device camera (CoolSNAP ES2) and captured into MetaVue imaging software v6.2. Images were taken every 30 s for 30–45 min with up to eight devices being imaged simultaneously. Tracking and velocity quantification were performed as described *(9).*

*Statistical Analysis:*

Statistical analysis was carried out using PRISM 6 (GraphPad). Mann–Whitney test (comparing two groups), Kruskal–Wallis test (when comparing to single group), and Gehan–Breslow–Wilcoxon test (for survival curve comparison) were used in this study as indicated in the figure legends. Individual p values are indicated in the figure with no data points excluded from statistical analysis. One representative experiment of at least three independent repeats are shown. For qPCR, each gene was normalized to a reference gene and compared with Holm-Sidak test (individual comparison with paired value) where each p value and DF was adjusted for respective multiple comparisons.

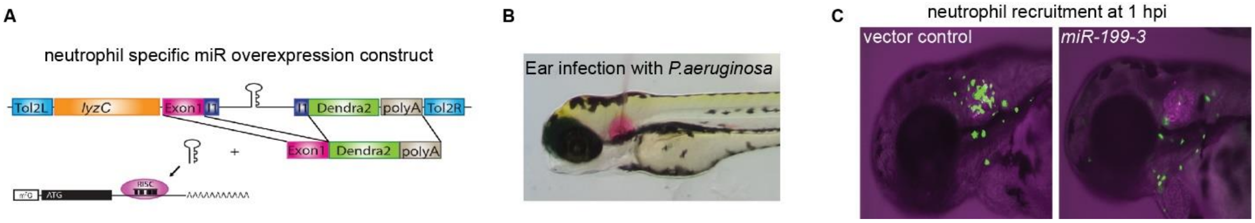


**Fig. S1. MicroRNA screen experimental design.**

(A) Each of the miR candidate gene is cloned in the intron of a green fluorescence reporter, driven by a neutrophil specific lyzC promoter. The plasmid is injected into 1-cell stage zebrafish embryos to induce expression of miR in a subset of neutrophils. (B) A representative picture of a bacterial infection which induces neutrophil recruitment to the otic vesicle. (C) Decreased neutrophil recruitment due to the overexpression *miR-199-3*. An inhibition of 50% neutrophil infiltrating into the ear indicates a positive hit.

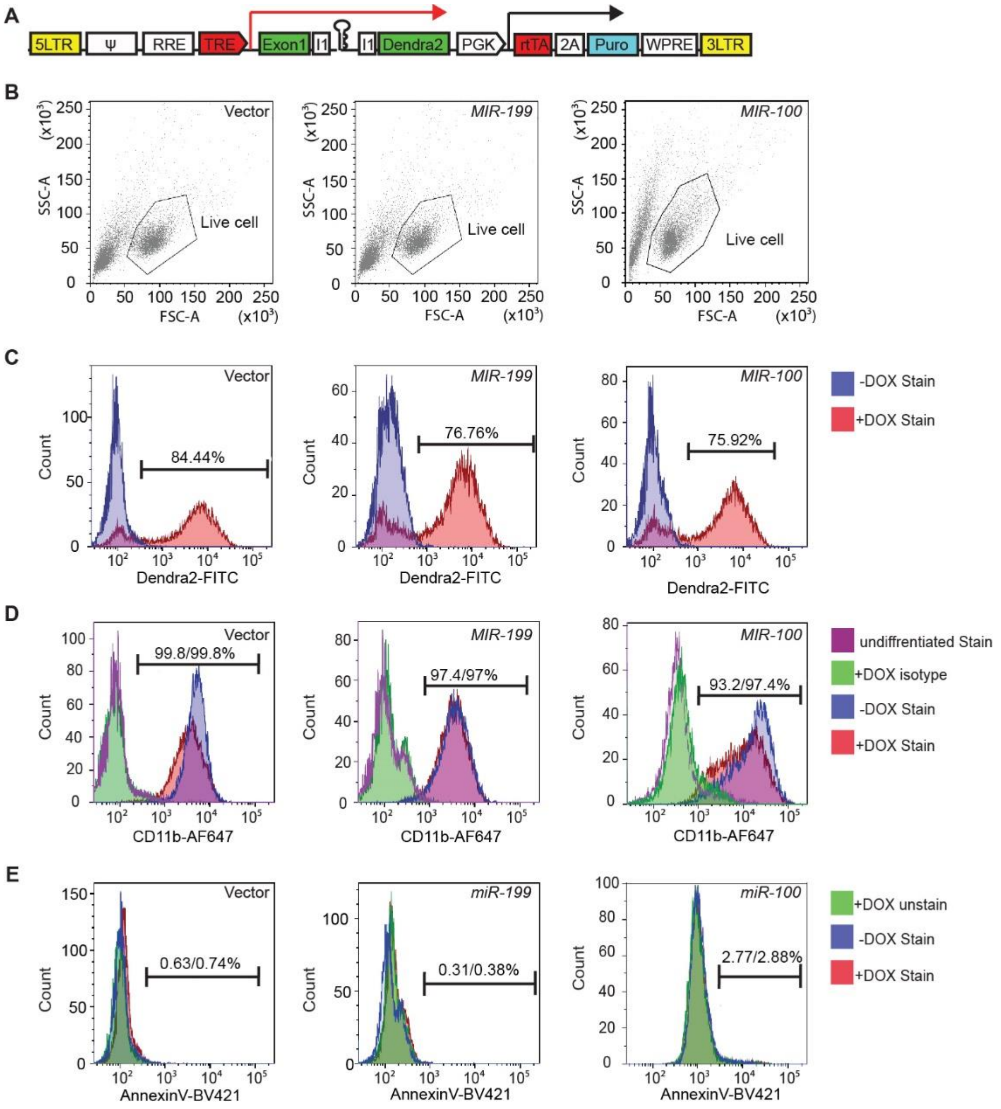


**Fig. S2. Characterization of HL-60 cell lines with miR overexpression induced after cell differentiation.**

(A) Construct for inducible miRNA expression. The miRNA and a dendra2 reporter (green) is under the control of TRE, Tetracyclin response element (red). The PGK promoter drives constitutive expression of puromycin resistance gene (cyan) and the rtTA, reverse tetracycline-controlled transactivator (red). (B) Cell populations gated for downstream cell profile analysis and transwell quantification. (C) Doxycycline mediated induction in the vector, *miR-199*, or *miR-100* expressing HL-60 lines. Cells without doxycycline mediated induction were used as a baseline. Percentage of cells with dendra2 level above the baseline are shown. (D) Surface CD11b levels in vector, *miR-199*, or *miR-100* expressing HL-60 cells. Undifferentiated cells or differentiated cells stained with an isotype control were used as a baseline. Percentage of cells with CD11b levels above the baseline with or without doxycycline mediated induction are shown. (E) Annexin V staining in the total cell population (excluding debris) of vector, *miR-199*, or *miR-100* expressing HL-60 cells. An unstained sample is used to determine the baseline.

Percentage of cells with AnnexinV levels above the baseline with or without doxycycline mediated induction are shown. (B-E) One representative experiment of three independent trials is shown.

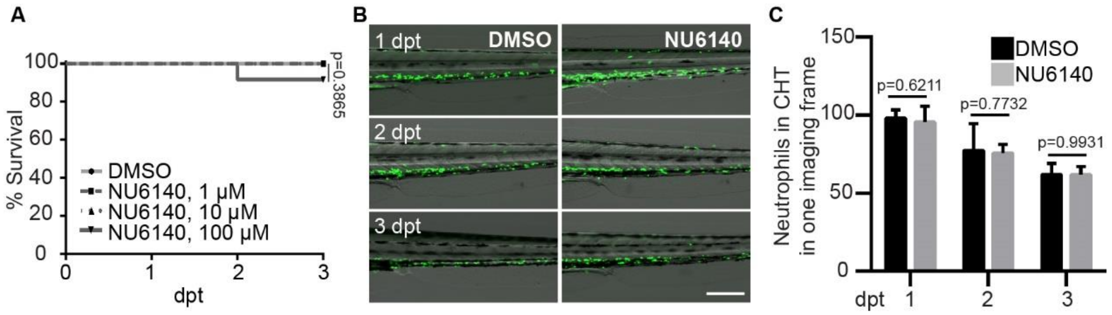


**Fig. S3. Cytotoxicity of CDK2 inhibitor NU6140.**

(A) Survival of 3 dpf zebrafish embryos treated with indicated doses of NU6140. Survival rates were tallied daily. One representative experiment of three independent experiments (n =20 each group) is shown, Gehan–Breslow–Wilcoxon test. (B, C) Representative images (B) and quantification (C) of neutrophils in the caudal hematopoietic tissue of the 3 dpf embryos from *Tg(lyzC:GFP)* treated with 100 μM of NU6140 for 3 days. Results are presented as mean ± s.d. of three individual trials, Kruskal–Wallis test.

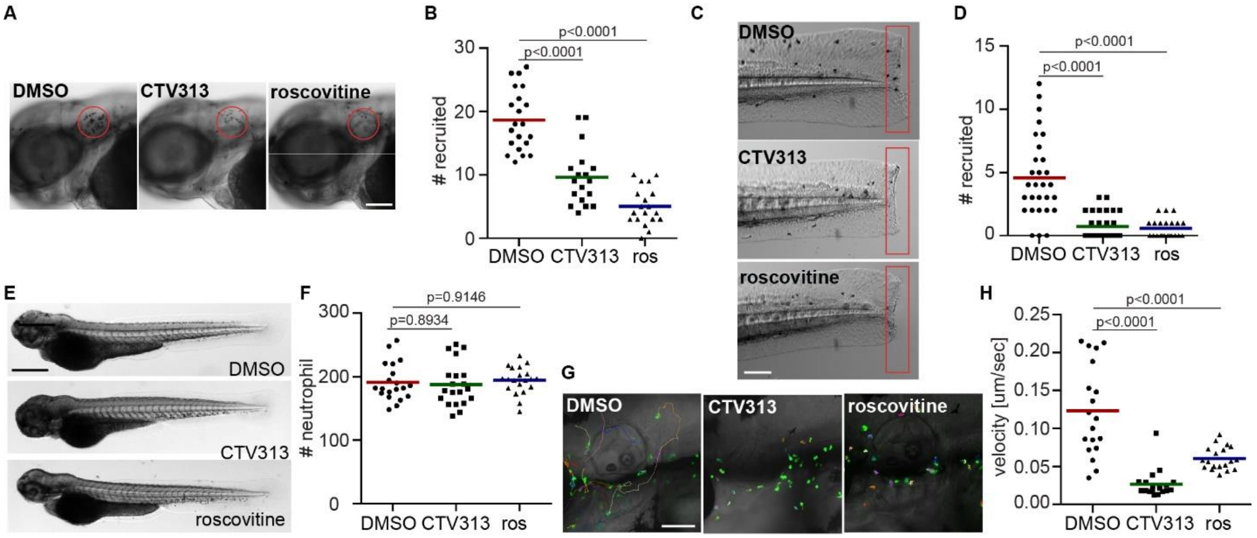


**Fig. S4. The CDK2 inhibitor CTV313 or a pan cdk inhibitor roscovitine reduces neutrophil migration.**

(A, B) Representative images (A) and quantification (B) of neutrophils recruited to infected ear in zebrafish larva treated with a cdk2 inhibitor (CTV313) or a pan-CDK inhibitor (roscovitine). Scale bar: 100 µm. (C, D) Representative images (C) and quantification (D) of neutrophils recruited to tail fin transection sites in zebrafish larva treated with CTV313 or roscovitine. Scale bar: 200 µm. (E, F) Representative images (E) and quantification (F) of total neutrophil number in zebrafish larva treated with CTV313 or roscovitine. Scale bar: 500 µm. (A-F) Assays were done with at least 3 individual founders with 3 biological repeats each containing 20 (for motility) or 25 (for neutrophil recruitment) fish per group. Result is presented as mean ± s.d., Mann–Whitney test. (G, H) Representative images (G), and quantification of velocity (H) of neutrophils in zebrafish larvae treated with CTV313 or Roscovitine. Scale bar 100 µm. 3 embryos each from three different founders were imaged and quantification of neutrophils in one representative movie is shown, Kruskal–Wallis test.

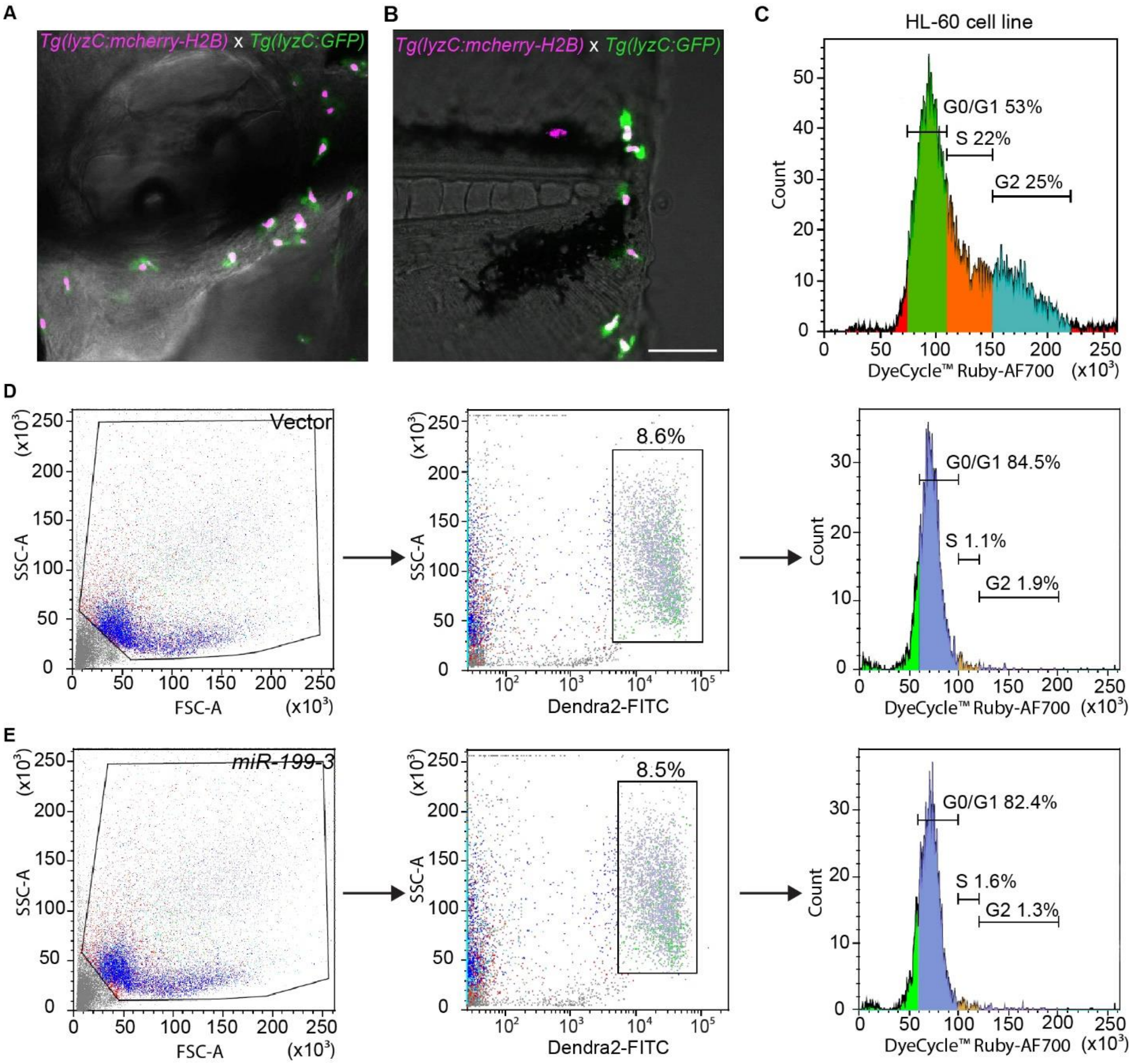


**Fig. S5. Cell cycle profiling of zebrafish neutrophils.**

(A) A representative image of 3 dpf embryos from *Tg(lyzC:mcherry-H2B)* (nucleus magenta label) crossed with *Tg(lyzC:GFP)* (cytosol green label) at 30 min post ear infection. One representative image of three independent experiments are shown. (B) A representative image of 3 dpf embryos from *Tg(lyzC:mcherry-H2B)* (nucleus red label) crossed with *Tg(lyzC:GFP)* (cytosol green label) at 30 min post tail wounding as described in (A). (C) Cell cycle profile of HL-60 cell line. Cells were separated into G1, S and G2 phased based on the fluorescence intensity of the cell cycle dye. (D) Cell cycle profile (right panel) of neutrophils in adult kidney marrow from the vector control line. Live cells were gated (left panel) and dendra+ cells were selected for analysis (middle panel). One representative experiment of three biological repeats are shown. (E) Cell cycle profile of neutrophils in adult kidney marrow from the *miR-199* line as described in (D).

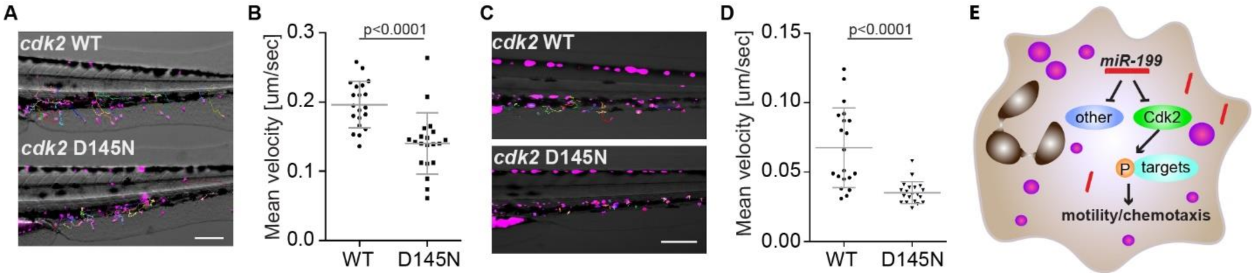


**Fig. S6. Cdk2 D145N inhibits neutrophil motility and chemotaxis and the working model.**

(A) Representative images and (B) quantification of velocity of neutrophils in 3 dpf zebrafish embryos from the WT or D145N line in a LTB4 bath. Scale bar: 200 µm. (C) Representative images and (D) quantification of velocity of neutrophil motility in 3 dpf zebrafish embryos from the WT or D145N line in the caudal hematopoietic tissue. Scale bar: 200 µm. Assays were performed with at least 3 individual founders with 3 biological repeats each containing at least 3 fish per group. Result is presented as mean ± s.d., Mann–Whitney test. (E) Working model of how *miR-199* inhibits neutrophil migration.

**Data file S1. Relative expression of miRs in sorted zebrafish neutrophils and apical keratinocytes and whole larva at 3 dpf.**

**Data file S2. Quantification of percentages of neutrophils recruited to the infected ear upon overexpression of selected miRNAs.**

**Data file S3. Quantification of numbers of neutrophils recruited to the infected ear and tail wounding sites in transgenic lines.**

**Data file S4. List of genes that are downregulated in the *miR-199* overexpressing neutrophils.**

**Data file S5. Pathway enrichment analysis of downregulated genes in *miR-199* overexpressing neutrophils.**

**Movie S1: Tracked movies of neutrophil motility lines in the head mesenchyme of the vector and the *miR-199*.**

The video shows the tracked baseline motility of neutrophils in 3 dpf vector and *miR-199-3* zebrafish lines. Videos were recorded for 30 min with 1 min interval. Representative videos from n = 3 independent experiments with 3 fish each group are shown. Scale bar: 100 µm.

**Movie S2: Tracked movie of neutrophil motility in the head mesenchyme treated with vehicle, NU6140, Aphidicolin+hydoxyurea, CTV313, or Roscovitine.**

The video shows the tracked baseline motility of neutrophils in 3 dpf zebrafish larvae treated with DMSO (vehicle control), NU6140 or Aphidicolin+hydoxyurea, CTV313 or roscovitine. Videos were recorded for 30 min with 1 min interval. Representative videos from n = 3 independent experiments with 3 fish each group are shown. Scale bar: 100 µm.

**Movie S3: Neutrophils are not dividing after recruited to regional inflammation sites.**

The video shows neutrophil recruitment to ear infection or tail wounding sites in zebrafish embryos at 3 dpf. Neutrophil cytosols were labeled green and nucleus labeled magenta. Videos were recorded for 120 min with 2 min interval. Representative videos from n = 3 independent experiments with 3 fish each group are shown. Scale bar: 50 µm.

**Movie S4: NU6140 inhibits chemotaxis of primary human neutrophil.**

Video and Tracks of primary human neutrophils chemotaxis treated with DMSO or NU6140 towards IL-8. Videos were recorded for 50 min with 30 sec interval. Representative videos from n = 4 separate trials are shown. Scale bar: 50 µm.

**Movie S5: Tracked movies of neutrophil motility in the head mesenchyme of the Cdk2 WT and D145N lines.**

The video shows the tracked baseline motility of neutrophils in 3 dpf Cdk2 WT and D145N zebrafish lines. Videos were recorded for 30 min with 1 min interval. Representative videos from n = 3 independent experiments with 3 fish each group are shown. Scale bar: 100 µm.

**Movie S6: Tracked movies of neutrophil recruitment to the fin in the Cdk2 WT and D145N lines bathed in LTB4.**

The video shows the tracked chemotaxis of neutrophils in 3 dpf Cdk2 WT and D145N zebrafish lines in LTB4 bath. Videos were recorded for 30 min with 1 min interval. Representative videos from n = 3 independent experiments with 3 fish each group are shown. Scale bar: 200 µm.

**Movie S7: Tracked movies of neutrophil motility in in the caudal hematopoietic tissue of the Cdk2 WT and D145N lines**

The video shows the tracked baseline motility of neutrophils in 3 dpf Cdk2 WT and D145N zebrafish lines. Videos were recorded for 30 min with 1 min interval. Representative videos from n = 3 independent experiments with 3 fish each group are shown. Scale bar: 200 µm.

**Movie S8: Morphology of neutrophils from the Cdk2 WT and D145N lines**

The video shows the morphology and stable actin distribution of neutrophils in 3 dpf Cdk2 WT and D145N zebrafish lines crossed with a *Tg(mpx:GFP-UtrCH)* line. Videos were recorded for 30 min with 1 min interval. Representative videos from n = 3 independent experiments with 3 fish each group are shown. Scale bar: 20 µm.
